# Traditional martial arts and shooting training have different effects on auditory fine structure processing ability——Evidence from behavioral tests and fMRI

**DOI:** 10.1101/2021.12.05.471289

**Authors:** Yu Ding, Keying Zhang, Chunmei Cao

**Affiliations:** Division of Sports Science and Physical Education, Tsinghua University, Beijing, 100084 China

**Keywords:** traditional martial arts, shooting, fine structure, fMRI

## Abstract

Explore the influence of traditional martial arts and shooting training on the ability of auditory temporal fine structure (TFS) processing. Twenty-five college students participated in the experiment, including 8 traditional martial arts practitioners, 8 high-level shooting athletes, and 9 control groups without any regular exercise habits. The BIC (break in interaural correlation) delay threshold and TFS1 test were used to evaluate the temporary storage capacity and sensitivity of TFS, respectively, and a fMRI test was performed after the test. The results found that the traditional martial arts group had stronger TFS sensitivity, while the shooting group had stronger TFS retention ability, and the performance of the behavioral test of the shooting group is related to the fALFF value of the brain area of interest. Traditional martial arts and shooting training have improved the ability of auditory information processing from different angles, diversified exercise habits will lead to the development of diversity in brain structure and function.

## 1. Introduction

Physical training can have a plastic effect on the brain and thus change cognitive functions(Rogge, Röder, Zech, & Hötting, 2018). Among many kinds of sports, the influence of traditional martial arts and shooting training on cognitive function has obvious characteristics. Due to the complexity of the movements and meanings of traditional martial arts, the brain and cognitive functions can be effectively exercised in the process of learning traditional martial arts(Fujiwara et al., 2019; Sai Srinivas, Vimalan, Padmanabhan, & Gulyás, 2021). In addition, some training in traditional martial arts, such as standing stance, also has similar effects with mindfulness(Fujiwara et al., 2019). Shooting events originated from hunting activities. From an evolutionary point of view, shooting has extremely high requirements on the brain’s sensory and motor control systems(Momi et al., 2021). Whether it is traditional martial arts or shooting training, it involves the improvement of information processing ability, and the improvement of this information processing ability is likely to cross sensory channels, such as the improvement of auditory information processing ability.

Before the sound enters the auditory cortex, it is first decomposed into narrowband signals in the cochlea, and these narrowband signals can be further divided into slow fluctuating envelope (ENV) components and fast fluctuating temporal fine structure (TFS). ENV and TFS play important roles in content recognition and spatial positioning, that is, ENV and TFS represent the “what” pathway and the “where” pathway in the auditory cortex, respectively(Smith, Delgutte, & Oxenham, 2002). Among them, the binaural integration of the TFS is the key to auditory spatial positioning and speech recognition in complex scenes(Moore, 2019). Because binaural information usually has a interaural interval (IAI), to effectively integrate binaural information, the auditory system needs to save the TFS of leading wave for a very short time. Therefore, binaural information integration involves not only the sensitivity of the TFS of the single ear, but also the ability to temporarily store the TFS. The TFS sensitivity can be measured by the “TFS1” method(Moore & Sek, 2009), and the temporary storage capacity of the TFS can be measured by the “BIC (break in interaural correlation) delay threshold” test(H. Li, Kong, Wu, & Li, 2013).

The processing of TFS in the brain reflects the accuracy of the conversion from sound signals to neural signals. The sensitivity and the ability of temporary storage of TFS have decreased significantly with aging(L. Li, Huang, Wu, Qi, & Schneider, 2009; Moore, Vickers, & Mehta, 2012), which may be one of the reasons for the decline in the cognitive ability and the speech recognition ability in complex scenarios of the elderly. This study examines the effects of traditional martial arts and shooting training on the sensitivity and retention of TFS, and provides evidence from behavioral measurements and brain imaging.

## 2. Methods

### 2.1 Participants

Twenty-five college students (17 males and 8 females; mean age = 22.60 ± 3.50) with normal hearing participated in this study, including 8 traditional martial arts practitioners, 8 elite shooting athletes, and 9 normal people without any regular exercise habits. Their pure-tone thresholds were no more than 20 dB hearing level (HL) between 0.125 and 8 kHz (ANSI-S3.6, 2004) and the threshold difference between the two ears in each frequency was less than 15 dB HL. All subjects signed an informed consent form and received a certain amount of remuneration. All subjects underwent the BIC delay threshold test and TFS1 test, and the martial arts group and shooting group underwent fMRI scans.

### 2.2 Apparatus

All participants were tested in a sound-attenuated room, environmental noise is less than 29 dB SPL. All the acoustic signals were calibrated by a sound-level meter (AUDit and System 824, Larson Davis, USA), and delivered using the Creative Sound Blaster (Sound Blaster X-Fi Surround 5.1 Pro, Creative Technology Ltd, Singapore), and presented to participants by headphones (HD 650, SENNHEISER, Germany).

### 2.3 Test for storage capacity of TFS

The BIC delay threshold test has been described in detail in the past research(H. Li et al., 2013; L. Li et al., 2009). Put it simply, in each trial, the subject will hear two noises, one of which has an irrelevant segment (the BIC segment) inserted in the noise. The task of the subject is to select the noise with the BIC segment. Gradually increase the IAI, and use the three-up one-down procedure(Levitt, 1971) to find the max IAI as the threshold.

Before the BIC test, all the participants became familiarized with binaurally presented noise either with or without the BIC. The IAI started from 0 ms, increased following three consecutive correct identifications of the presentation containing the BIC, and decreased following one incorrect identification. The initial step size of changing the IAI was 16 ms, which was altered by a factor of 0.5 with each reversal of direction until the minimum size of 1 ms was reached. Visual feedback was given after each trial to indicate whether the identification was correct or not. A test session was terminated following 10 reversals in direction, and the longest IAI for a session was defined as the mean IAI for the last 6 reversals. The average over the three repeated-test sessions was used as the longest IAI for each participant.

### 2.4 Test for sensitivity of TFS

In the testing stages of TFS1, this study uses the special software package published by moore on the Internet(https://www.psychol.cam.ac.uk/hearing)(A. P. Sęk & Moore, 2020). Most of the parameters use the default settings(A. Sęk & Moore, 2012). The fundamental frequency of the TFS1 test is 200 Hz, the center frequency is 1800 Hz, the signal sound intensity is 60 dB SPL, the noise sound intensity is 45 dB SPL, and the initial change frequency is 100 Hz. All the results are automatically output by the software after the test.

### 2.5 MRI scan

MRI images were acquired on a 3 Tesla Philips Achieva scanner using a 32-channel head coil at the Center for Biomedical Imaging Research of Tsinghua University. During each scan sequence, subjects were told to relax and remain awake with their eyes closed. Headphones and foam were used for added hearing protection against the machine noise. Eight minutes of Echo Planar Imaging(EPI) sequence was used to acquire Blood oxygen level-dependent (BOLD) data and the parameters were as follows: TE = 30 ms; TR = 2 s; flip angle = 90 degree; slices = 37 sagittal slices; slice thickness = 3 mm; voxel size = 2.87 × 2.87 × 3.50 mm3; image matrix = 80 × 80; FOV = 230 × 230 mm, slices were obtained starting from the bottom of the cerebellum to the top of the head. Structural T1-weighted images were also acquired with the parameters of: flip angle = 8 degree; slices = 180 sagittal slices; voxel size =1 × 1 × 1 mm3; FOV = 230 × 230 mm, slices were obtained starting from the right side to the left side of the brain.

### 2.6 VBM analysis

VBM analysis was used to analyze the gray matter volume plasticity and was performed using VBM8 toolbox (http://dbm.neuro.uni-jena.de/wordpress/vbm/download/). First, the high-resolution images were normalized to standard space using DARTEL method. Then, the normalized images were segmented into different tissues white matter, and cerebrospinal fluid. Finally, the gray matter images were normalized to the MNI template and then smoothed with a Gaussian kernel of 4 × 4 × 4 mm3 of FWHM.

### 2.7 Resting state pretreatment and fALFF calculation

Preprocessing was carried out DPARSF (http://rfmri.org/DPARSF) package based on MATLAB in regular order and with default parameters. First, five time points were removed to exclude the influence of the instability of the initial MRI signal. Slice timing and head motion correction were then carried out to correct the influence of within scan acquisition time difference and the subject’s head movements. Subjects with a head motion of >2 mm in any plane or rotation >2° in any direction were excluded. Realigned images were then spatially normalized to the high-resolution T1-weighted images in the standard Montreal Neurological Institute (MNI) space and were resampled to a voxel size of 3 × 3 × 3 mm3 for the sake of group comparison. After that, covariates were removed including regression of linear trend, and 6-Friston head motion. Finally, spatial smoothing was then conducted with a Gaussian kernel of 4 × 4 × 4 mm3 of full width at half maximum (FWHM). In this research, an fALFF (Zou et al., 2008) map was calculated by DPARSF for each subject using the preprocessed data. The Fisher z transformation was then performed on the fALFF graph for statistical analysis.

## 3. Results

### 3.1 behavioral performance of participants

Figure 1 shows the behavioral performance of different groups for BIC delay threshold test, the ANOVA test showed that the main effects of different groups were significant, F(2,22) = 4.560, *p* = 0.022, *η*_*p*_^*2*^, and the post-test found that the shooting group was significantly higher than the normal control group, *p* = 0.022. For TFS1 test, the ANOVA test showed that the main effects of different groups were significant, F(2,22) = 4.051, *p* = 0.032, *η*_*p*_^*2*^, and the post-test found that the martial arts group was significantly lower than the normal control group, *p* = 0.032. Note that for the BIC delay threshold test, the higher the score, the longer the retention time. For the TFS1 test, the lower the score, the better the sensitivity. Therefore, martial arts and shooting improve the ability of TFS processing at different angles.

**Figure 1.**
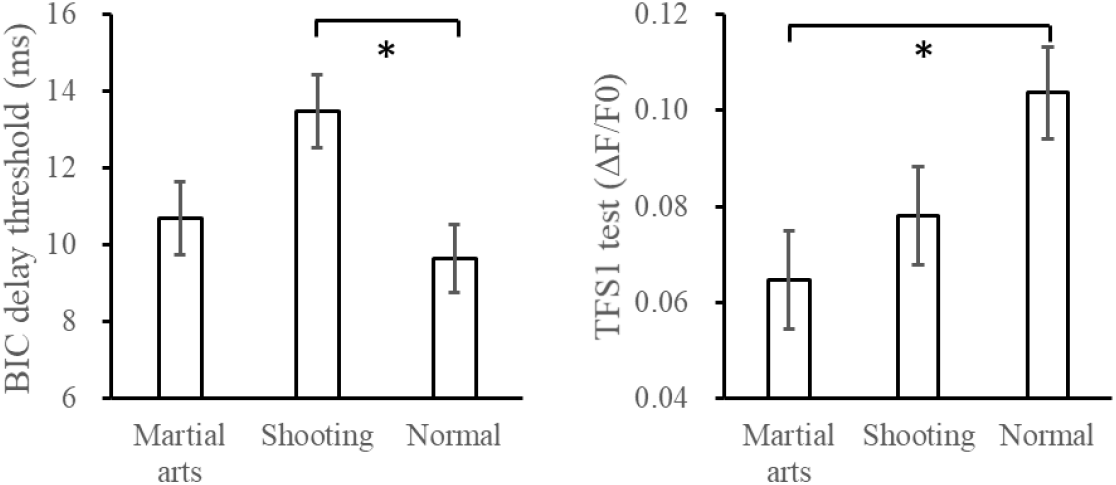
behavioral performance of different groups

### 3.2 VBM and fALFF

The following tables show the results of VBM and fALFF. Statistical maps were assessed at an uncorrected threshold of p < 0.01; the two clusters were significant at p < 0.01, familywise-error-corrected, at the cluster level. Coordinates are given in Montreal Neurological Institute (MNI) space. Peak T > 0 means aerobic group > anaerobic group, while peak T < 0 means aerobic group < anaerobic group.

**table 1.**
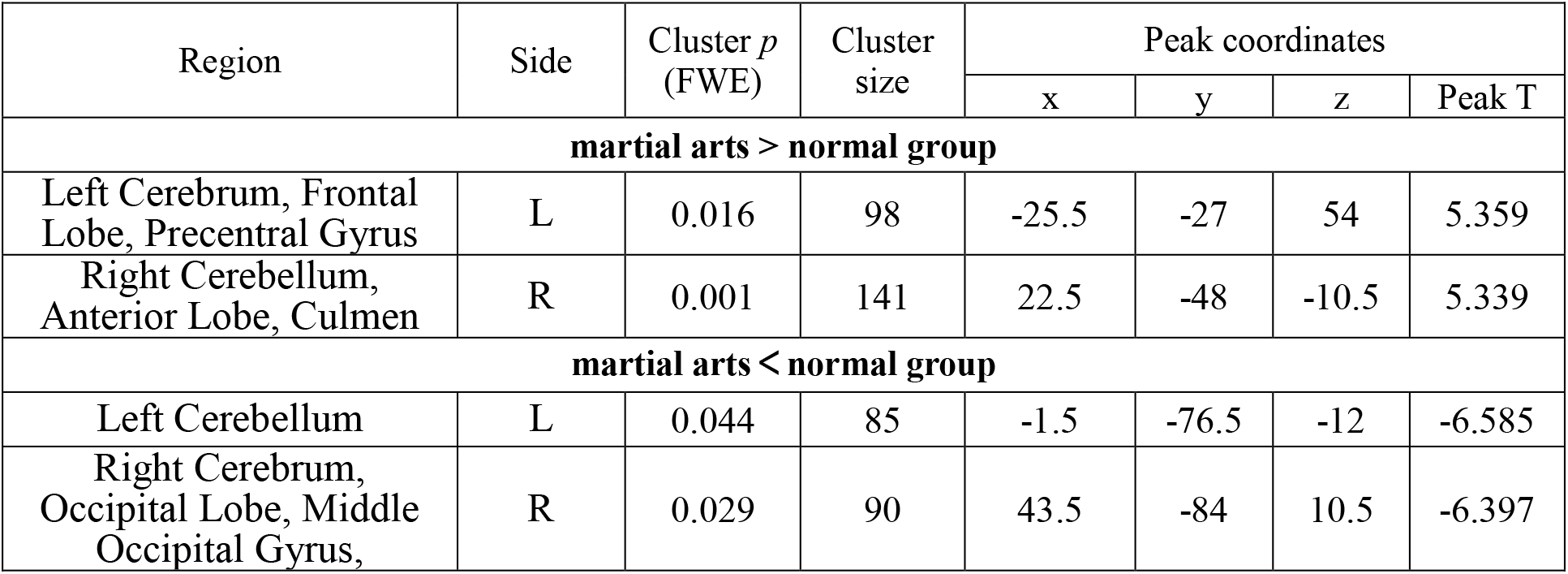
VBM: **martial arts vs normal group**

**table 2.**
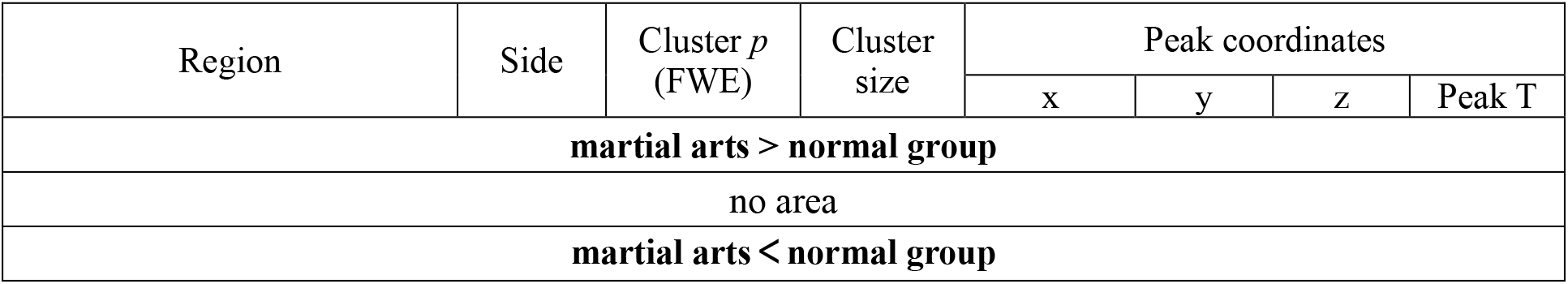
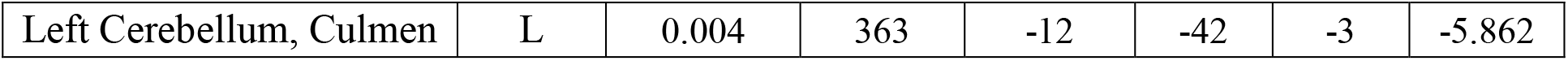
fALFF: **martial arts vs normal group**

**table 3.** VBM **shooting vs normal group**

| Region | Side | Cluster *p*<br>(FWE) | Cluster<br>size | Peak coordinates |  |  |  |
| --- | --- | --- | --- | --- | --- | --- | --- |
|  |  |  |  | x | y | z | Peak T |
| shooting> normal group |  |  |  |  |  |  |  |
| Left Cerebellum,<br>Anterior Lobe, Culmen | L | 0.000 | 165 | -43.5 | -48 | -25.5 | 6.086 |
| shooting< normal group |  |  |  |  |  |  |  |
| no area |  |  |  |  |  |  |  |

**table 4.** fALFF **shooting vs normal group**

| Region | Side | Cluster $p$<br>(FWE) | Cluster<br>size | Peak coordinates | | | |
| --- | --- | --- | --- | --- | --- | --- | --- |
|  |  |  |  | x | y | z | Peak T |
| shooting> normal group |  |  |  |  |  |  |  |
| Right Cerebrum, Frontal<br>Lobe, Middle Frontal<br>Gyrus | R | 0.000 | 920 | 45 | 27 | 48 | 4.643 |
| Right Cerebrum, Parietal<br>Lobe, Inferior Parietal<br>Lobule | L | 0.002 | 418 | 48 | -51 | 48 | 4.490 |
| shooting< normal group |  |  |  |  |  |  |  |
| Left Cerebellum,<br>Posterior Lobe, Inferior<br>Semi-Lunar Lobule | L | 0.003 | 388 | -6 | -69 | -42 | -4.756 |
| Right Cerebellum,<br>Posterior Lobe, Pyramis | R | 0.010 | 313 | 12 | -72 | -24 | -4.521 |

**table 5.** VBM **martial arts vs shooting**

| Region | Side | Cluster *p*<br>(FWE) | Cluster<br>size | Peak coordinates |  |  |  |
| --- | --- | --- | --- | --- | --- | --- | --- |
|  |  |  |  | x | y | z | Peak T |
| martial arts > shooting |  |  |  |  |  |  |  |
| Right Cerebellum,<br>Posterior Lobe, Declive | R | 0.000 | 279 | 27 | -60 | -18 | 6.195 |
| martial arts < shooting |  |  |  |  |  |  |  |
|  | L | 0.001 | 135 | 13.5 | -24 | 76.5 | -7.257 |
| Right Cerebrum, Frontal<br>Lobe, Precentral Gyrus, | R | 0.021 | 94 | 37.5 | -19.5 | 66 | -5.120 |
| Left Cerebrum, Parietal<br>Lobe, Sub-Gyral | L | 0.011 | 103 | -22.5 | -42 | 57 | -5.057 |
| Left Cerebrum, Sub-<br>lobar, Extra-Nuclear | L | 0.017 | 97 | -12 | 15 | -9 | -4.181 |

**table 6.** fALFF **martial arts vs shooting**

| Region | Side | Cluster *p*<br>(FWE) | Cluster<br>size | Peak coordinates |  |  |  |
| --- | --- | --- | --- | --- | --- | --- | --- |
|  |  |  |  | x | y | z | Peak T |
| martial arts > shooting |  |  |  |  |  |  |  |
| Left Cerebrum, Frontal | L | 0.000 | 1469 | -48 | 9 | 51 | 4.760 |
| Lobe, Middle Frontal Gyrus |  |  |  |  |  |  |  |
| Left Cerebrum, Parietal Lobe, Superior Parietal Lobule, Gray Matter | L | 0.030 | 274 | -30 | -57 | 63 | 3.919 |
| **martial arts < shooting** |  |  |  |  |  |  |  |
| Right Cerebellum, | R | 0.010 | 341 | 27 | -69 | -54 | -2.992 |

### 3.3 Correlation between behavioral performance and fALFF

Figure 2 shows the correlation between behavioral performance and fALFF in shooting group. BIC delay threshold is significantly related to fALFF in middle frontal gyrus (*r* = 0.728, *p* = 0.040) and fusiform gyrus (*r* = -0.774, *p* = 0.024), while TFS1 is significantly related to fALFF in fusiform gyrus (*r* = 0.847, *p* = 0.008) and inferior tempor gyrus (*r* = -0.756, *p* = 0.030). No significant correlation was found in the martial arts group (all p > 0.05).

**Figure 2.**
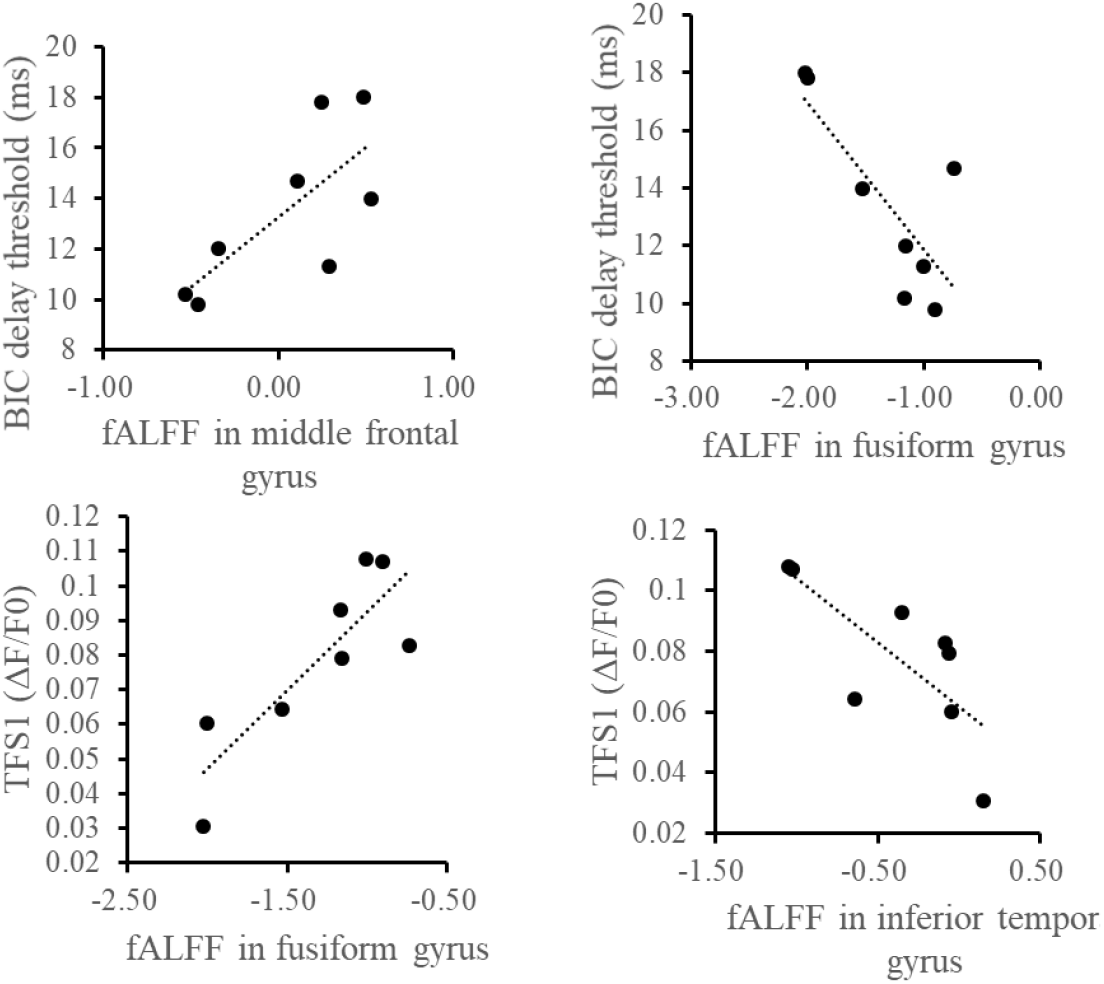
Correlation between behavioral performance and fALFF in shooting group

## 4. Discussion

From the perspective of behavioral indicators, traditional martial arts and shooting training have different effects on auditory information processing ability. Specifically, compared with the non-exercise control group, subjects who practice traditional martial arts have better TFS sensitivity, while the shooter has better TFS retention ability. The appearance of such behavioral characteristics may be related to the training characteristics of these two sports. Traditional martial arts training often requires a momentary perception and reaction. Even one’s routine training often requires setting up an imaginary enemy and conducting imaginary training for avoidance and attack. It may be that these trainings increase the sensitivity of sensory information, and the increase in TFS sensitivity allows trainers to more keenly judge the spatial position of the opponent’s attack based on auditory information. This bottom-up information processing is very rapid. There are related concepts in many traditional martial arts, such as “Listening to the Bridge” in Wing Chun and “Listening Power” in the Tai Chi Push Hands. In the shooting training process, due to the target’s movement, air vibration, heart rate and breathing, and the regular shaking of the arms and torso, the shooter often needs to make a prediction about the timing of shooting based on the information obtained. Therefore, shooting is not only a sport based on spatial information, but also a sport based on time information. This anticipation of delay may train the subjects’ ability to maintain the fine structure over time.

Combined with the results of brain imaging, the behavioral test scores of shooting training have a higher correlation with the fALFF value of the area of interest, while no similar correlations can be seen in the martial arts group. There are many possible explanations for this. For example, the level of shooting training is easier to define. All shooting team members in this study are high-level athletes. The level of traditional martial arts is more difficult to define. Because traditional martial arts include many different schools and training methods, it is difficult to find an authoritative standard for evaluating ability. Although the behavioral results of this study found that traditional martial arts practitioners have stronger sensitivity to TFS, in fact their brain and cognitive functions are diversified. It is precisely because traditional martial arts are diverse, their diversified training methods and connotations have led to the diversified development of the brain and cognitive functions of the practitioners. Future research can conduct more detailed research from the perspective of specific training details.

There are two possible explanations for all the results of this study. One is that traditional martial arts and shooting training lead to plastic changes in brain structure and function, and further affect behavioral indicators. Another possible explanation is that due to genetic diversity, people with different behavioral abilities are more inclined to choose other sports such as traditional martial arts or shooting. Future longitudinal studies can obtain clearer data and conclusions through tracking or intervention.

## Acknowledgements

This work was supported by the National Defense Science and Technology Project (1716312ZT002081) and “Shuimu Tsinghua Scholar” Project (2020SM055). There is no conflict of interest between all authors.

## Notes

### Competing Interest Statement

The authors have declared no competing interest.

